# Obesity status and its relative factors of captive Asian elephants (*Elephas maximus*) in China based on body condition assessment

**DOI:** 10.1101/2023.11.25.568637

**Authors:** Yinpu Tang, Ting Jia, Fangyi Zhou, Liang Wang, Li Zhang

## Abstract

Obesity is a common health problem in captive wildlife. Since the obesity status of captive Asian elephants (*Elephas maximus*) in China has not previously been investigated, we recorded seven relevant variables (sex, age, daily feed supply, proportion of high-calorie feed, outdoor enclosure area, outdoor time, and foot disorder) related to obesity in 204 captive Asian elephants through field investigation of 43 elephant-raising facilities. Assessment of obesity was based on visual body condition scoring for each individual. It revealed that obesity was prevalent for captive Asian elephants (especially elephants in zoos) in China. Over 70% of captive Asian elephants in China were overweight or in obesity to various degrees. Statistical analysis showed that for elephants in zoos, insufficient outdoor time might be the primary potential cause of obesity. We suggested facilities to extend the outdoor time and control the supply of high-calorie feed (e.g., fruits, vegetables, pellets, etc.) for captive elephants, thereby alleviating obesity through increased exercise and a suitable energy intake. Moreover, all facilities should implement positive reinforcement training to facilitate regular physical examinations, including foot health checks and blood sampling. This training would improve the ability to collect more precise information relating to elephant health, obesity, and the evaluation of animal welfare.

## Introduction

Asian elephants (*Elephas maximus*), the largest terrestrial animal in Eurasia, are listed as Endangered (EN) on the IUCN Red List and CITES Appendix I. In the wild, Asian elephants are distributed in 13 countries in South and Southeast Asia, with a total population of 48,323 to 51,680 (Menon & Tiwari 2019). Approximately 16,000 Asian elephants live in captivity (e.g., zoos, wildlife sanctuaries, etc.) or semi-captivity (e.g., timber enterprises, in private, etc.) and are mostly used for transport or as draught animals, only about 8.75% (< 1,400) living in zoos (Clubb & Mason 2002; Robinson *et al* 2012; Brown 2019).

Many studies have shown that captive elephants are prone to a variety of health problems, leading to a significant reduction in lifespan. Meanwhile, due to the low reproductive rate, zoo-elephant populations are not self-sustaining (Wiese 2000; Clubb & Mason 2002; Clubb *et al* 2008; Thitaram 2012). Consequently, the state of the reproductive health of captive Asian elephants and influential factors have always been the focus of breeding management and care (Schmidt 1982; Schaftenaar & Hildebrandt 2006; Kumar *et al* 2014; Pushpakumara *et al* 2016).

However, compared to physical injury or infectious disease, obesity, as a common health problem of wild animals in captivity, rarely received sufficient attention from caregivers (Schwitzer & Kaumanns 2001; Klimentidis *et al* 2011). In general, from the perspective of energy budget, an organism’s obesity status mostly depends on the nutritional intake and amount of exercise (Swinburn *et al* 2004; Shaw *et al* 2006). Therefore, an unsuitable diet and insufficient exercise tend to cause obesity in captive Asian elephants (Fowler 2008). According to a survey by the Association of Zoos and Aquariums (AZA) of the USA, among the 108 captive Asian elephants in 65 North American zoos, 75% of the females and 65% of the males are overweight or in obesity to various degrees (Morfeld *et al* 2016). Studies on captive Asian elephants in European and American zoos as well as Asian elephants engaged in Thai tourism suggest that obesity may indicate metabolic disorders in the body, which in turn are highly correlated with low fertility in both sexes (Taylor & Poole, 1998; Norkaew *et al* 2018; Norkaew *et al* 2019). In contrast, during a study of free-ranging Asian elephants in Sri Lanka, it was observed that appropriate body condition and sufficient physical exercise contribute to the breeding success of females. Moreover, relative body weight (kg/cm at shoulder) in elephants was positively correlated with reproductive output (Dastig 2002).

The history of raising Asian elephants in Chinese zoos can be traced back to 1907 (Yang *et al* 2002). Subsequently, the first instance of breeding Asian elephants in human cares took place in 1966 (Zhang 2018). Since the 1980s, China has begun to import a large number of Asian elephants from abroad (Zhang 2018). The population of captive Asian elephants has increased rapidly with the successful breeding of Asian elephants in various facilities. However, compared to European countries and the United States, China still has many shortcomings regarding the research and management of Asian elephants in captivity. One example of a knowledge gap is understanding obesity and its harm to captive Asian elephants. This study aimed to explore the relationship between potential related factors and the obesity status of captive Asian elephants in China. In addition, this study further summarized and discussed the main challenges faced by captive Asian elephants in China, and provided operational suggestions and physical conditions to improve their welfare status.

## Materials and methods

### Ethics statement

All the data collected in this study were for scientific purposes. Both Endangered Species Scientific Commission, P.R.China (ESSC, P.R.C) and Chinese Association of Zoological Gardens (CAZG) reviewed all the procedures and approved permits for this study conducted in the elephant-raising facilities below. Our study adopted observation-based methods without direct contact with animals or using tissue samples, and approval from an Institutional Animal Care and Use Committee or equivalent animal ethics committee was therefore not required.

### Animals

We recorded a total of 204 captive Asian elephants (♂:♀=88:116, about 64% of the living captive Asian elephant population in China) from January 2017 to April 2019 by on-site investigation of 43 facilities (including 42 zoos and Wild Elephant Valley in Xishuangbanna, see in “***Study areas***”). By interviewing staff members (veterinarians, curators, and caregivers) and consulting animal archives during our investigation, we recorded the sex and age of each elephant. However, the age of 31 individuals (♂:♀=13:18) was unknown (**Figure 1**).

**Figure 1.**
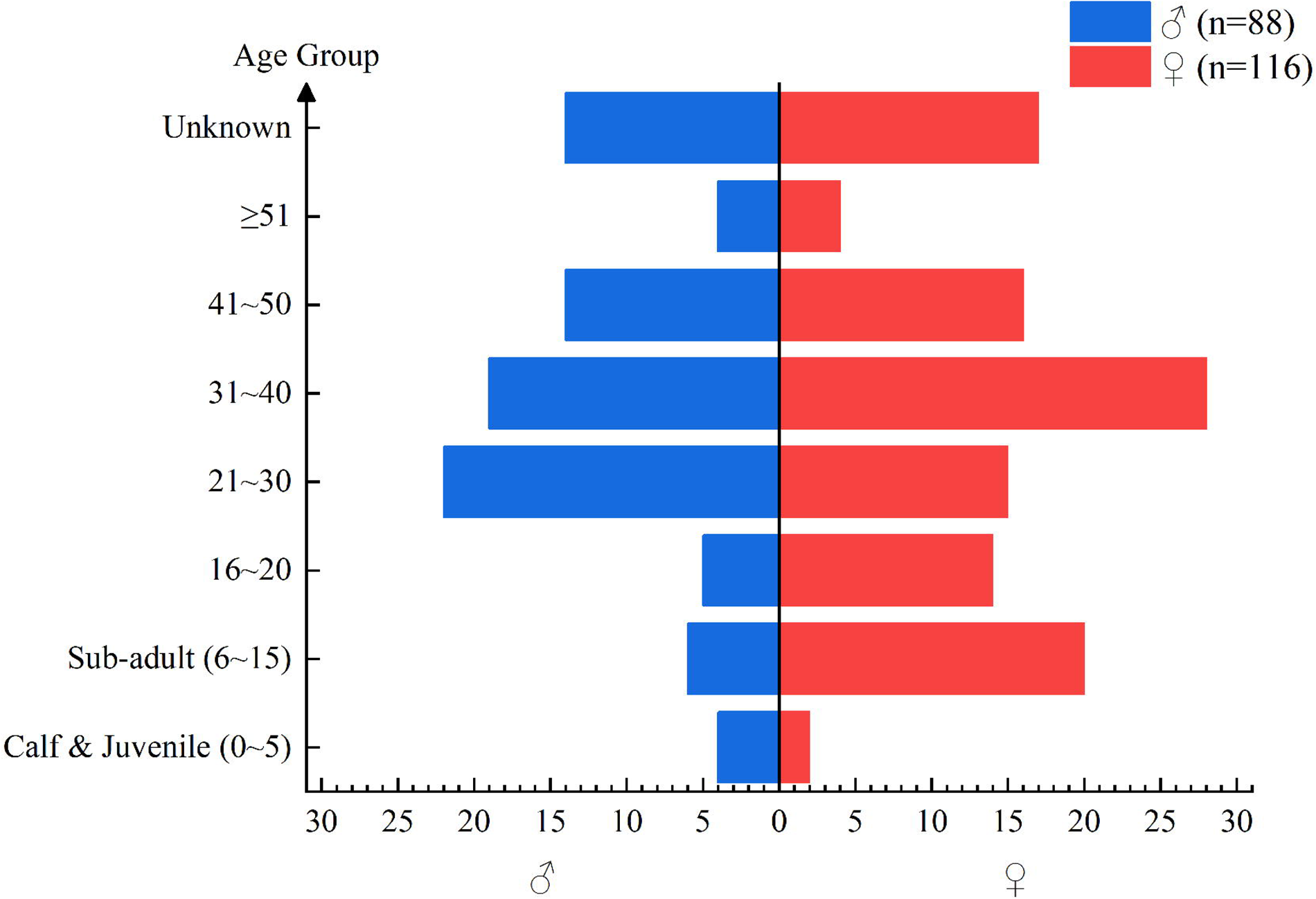
Age structure of 224 captive Asian elephants recorded from zoos and Wild Elephant Valley in China from January 2017 to April 2019. Classification of age groups was referred to the criteria in Arivazhagan & Sukumar (2008).

### Study areas

#### Zoos

A total of 42 zoos were visited on-site in this study, distributed in 25 provinces, municipalities, and autonomous regions. Each zoo usually opens at 9:00 and closes at 17:00 every day. When the weather is warm, elephants are allowed on display in outdoor enclosures during opening hours and return indoors when zoos are closed. Due to the lack of outdoor heating equipment, most zoos (especially those located in Northern China) no longer exhibit elephants outdoors when the temperature turns cold (e.g., ≤15°C) every year, and resume outdoor exhibitions until when it gets warmer next year. Caregivers provide each elephant with a certain amount of feed every day, which is divided into two meals in the morning and afternoon.

#### Wild Elephant Valley

The Wild Elephant Valley (100°51’33.2172’’E, 22°10’39.3636’’N) is located in Xishuangbanna National Nature Reserve, Yunnan province. As a tourist attraction and an elephant rescue center, it has a mixed population of imported elephants, which are trained for performance, as well as captive-born and rescued elephants. Except for performance time (about two hours every day), elephants here are allowed to forage and roam into a nearby natural forest region of 340 mu (approximately 226,666.67 m^2^, including 8,000 m^2^ of surface water and 2,800 m^2^ of local villages) during the day accompanied by their mahouts. Elephants here are provided with four tons of fresh grasses (mainly elephant grasses *Cenchrus purpureus*, sometimes with rye grasses *Leymus chinensis* and corn stalks, and 100 kilograms of seasonal fruits in total every day. During feeding hours, elephants usually form small groups (four to ten individuals) by themselves. Members of the same group eat the same pile of feed together, separating from other groups.

### Obesity status assessment – body condition scoring

Due to the lack of training and equipment in most study areas, we could hardly weigh their elephants directly. Consequently, we used the result of visual body condition assessment, a widely used method to evaluate the overall condition of animals, to represent the obesity status of elephants. This method was first adopted to assess the physical condition of large livestock such as cows and horses and has since been extended to wildlife, both in natural habitats and captivity (Carroll & Huntington 1988; Edmonson *et al* 1989; Schiffmann *et al* 2017; Sun *et al* 2021). In this study, we assessed each elephant’s body condition score (BCS) based on the criteria by Fernando *et al* (2009) (**Table 1**) and hypothesized that a higher BCS implied higher obesity status of an elephant (Chusyd *et al* 2019).

**Table 1.** Criteria of visual body condition scoring (BCS) used in this study to evaluate Asian elephants (*Elephas maximus*) for obesity (from Fernando et al 2009)

| Score | Characters |
| --- | --- |
| 1 | All ribs (shoulder to pelvis) visible, some ribs prominent (spaces in between sunken in) |
| 3 | Some ribs visible (spaces in between not sunken in), shoulder and pelvic girdles prominent |
| 5 | Ribs not visible, shoulder and pelvic girdles visible |
| 7 | Backbone visible as a ridge, shoulder and pelvic girdles not visible |
| 9 | Back rounded, thick rolls of fat under neck |

\* When body condition of an elephant is intermediate between two criteria, an intermediate point score (i.e. 2, 4, 6, 8 points) should be assigned. To accommodate the possibility of greater variation, the range of scores extend from 0 to 10.

Photographs and videos of elephants from various angles were taken during the investigation of each facility. Selected images (photographs and video screenshots) for body condition assessment had to fulfill the following requirements: (i) clear, identifiable individual; (ii) standing or moderate walking body position of each elephant to allow reliable assessment; (iii) sufficient recognition of prominent bone structures (skull, pectoral girdles in shoulders, vertebral column, ribs, pelvic girdles, and backbone); and (iv) adequate resolution for recognition of the generic wrinkles on the skin surface. Finally, (v) distinct patterns of shade or masses of hay, straw, and other substrates on the back of the elephant could make any assessment impossible. Likewise, bright lateral rays of sunlight could also reduce the image contrast and disturb the evaluation. Such documents were excluded from the study. All the elephant images were scored by the first author. Each individual was scored twice based on images both on the left and right sides, and an average score of the same individual on both sides was rounded up as the BCS of this individual. Here we considered that individuals with BCS ≥ 7 were in obesity.

### Potential relative factors of BCS

Including sex and age, we investigated seven potential relative factors of BCS (**Table 2**) in total, five of which were associated with diet and exercise.

**Table 2.**
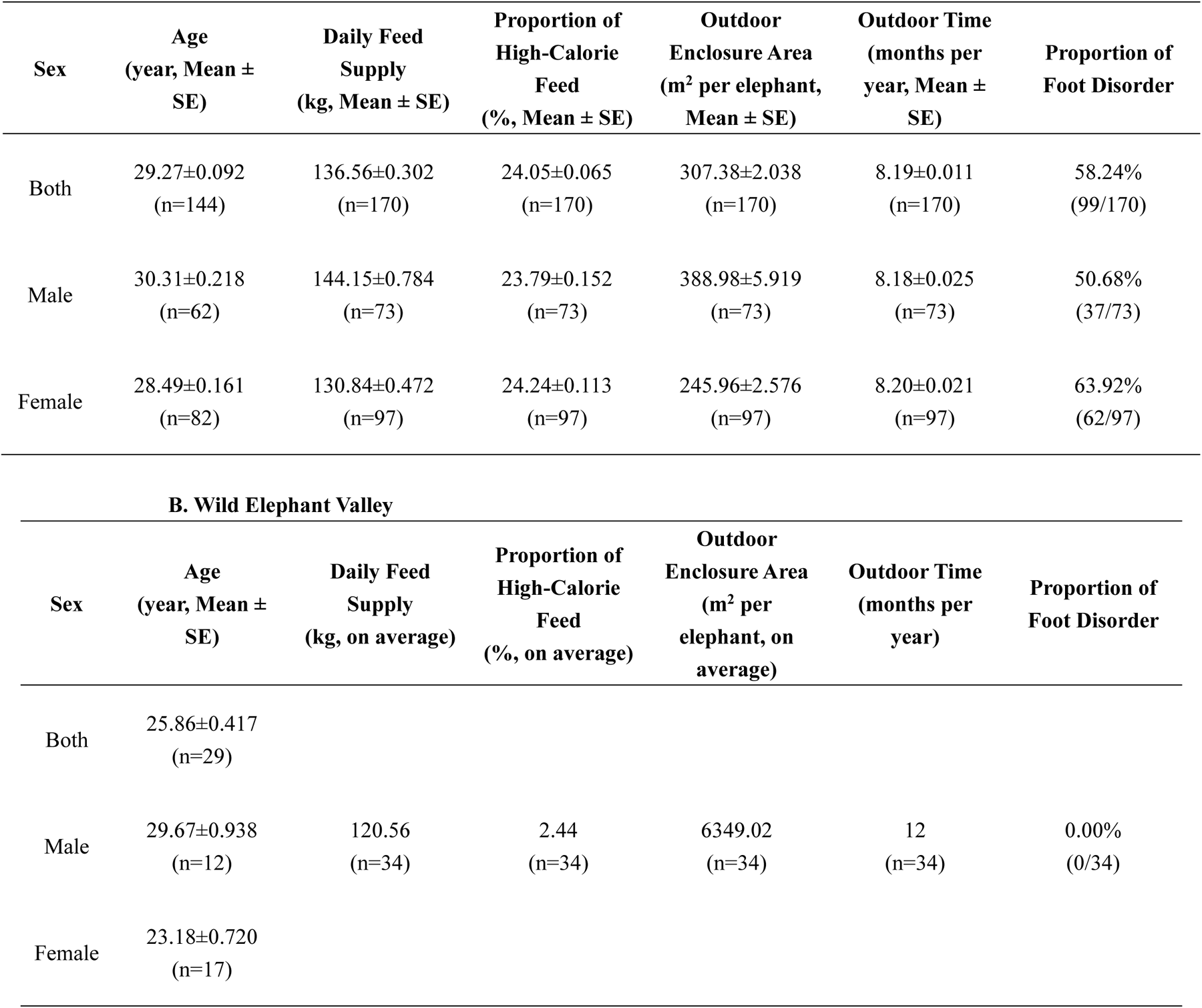
Overview of seven potential relative factors of BCS for elephants in different facilities. (A) Zoos; (B) Wild Elephant Valley

#### Potential relative factors of BCS associated with diet

There were two potential relative factors of BCS associated with diet: daily feed supply (kg) and the proportion of high-calorie feed (%).

Our investigation showed that for all 43 facilities, the feed of elephants was mainly divided into three categories: (i) herb forage, including various grasses and hays (e.g., Chinese rye grass, Sudan grass, medick, straw, etc.); (ii) juicy fodder (seasonal fruits and vegetables); and (iii) pellet feed, the main ingredients of which were corn kernels, wheat bran, soybean meal. Here we defined “high-calorie feed” as the sum of juicy fodder and pellet feed. For elephants in zoos, we collected the weight of three categories of feed above daily supplied for each elephant by interviewing staff members and consulting feeding documentation, then calculated the daily feed supply and the proportion of high-calorie feed of each elephant (**Table 2A**). For elephants in the Wild Elephant Valley, since it was challenging to ensure the exact daily feed supply for each elephant, we hypothesized that its daily total feed supply was distributed equally to each elephant (**Table 2B**). Here we hypothesized that both a larger daily feed supply and a higher proportion of high-calorie feed could associate with higher BCS in an elephant.

#### Potential relative factors of BCS associated with exercise

There were three potential relative factors of BCS associated with exercise: outdoor enclosure area (m^2^ per elephant), outdoor time (months per year), and foot disorder.

To avoid unnecessary stress on the elephants, we refrained from direct contact with them during our investigation. Therefore, it was challenging to calculate precise data on the exercise amount of elephants by recording walking distances through activity-tracking bracelets (Chusyd *et al* 2021) or measuring elephants’ oxygen consumption through wearable air-collecting apparatus (Langman *et al* 1995). However, we hypothesized that compared to narrow indoor enclosures, larger outdoor enclosures with longer outdoor time could provide elephants with more space to move and display their natural behaviors, thereby increasing their amount of exercise and energy consumption. Additionally, foot disorders could affect elephants’ mobility, which would lead to a decrease in their amount of exercise and energy consumption.

For each facility, we investigated its outdoor enclosure for elephants (**Table 2**). The outdoor enclosure area was measured by selecting the corresponding region on the map in Google Earth Pro 7.3.2.5766. It was noted that local villages and surface waters were excluded from the calculation when determining the outdoor enclosure areas for elephants in the Wild Elephant Valley. The outdoor time for elephants in zoos was recorded by interviewing staff members and consulting monthly exhibiting documentation. Meanwhile, for each elephant, we took close-up shots of its four feet and recorded whether it had one or more of the following visible foot disorders: (i) overgrown nails; (ii) nail cracks; (iii) overgrown cuticles, and (iv) joint deformation (**Figure 2**, **Table 2**). If any of these four visible disorders were not observed in each foot of an elephant, then we considered that this elephant’s feet were in healthy appearance (**Figure 2A**).

**Figure 2.**
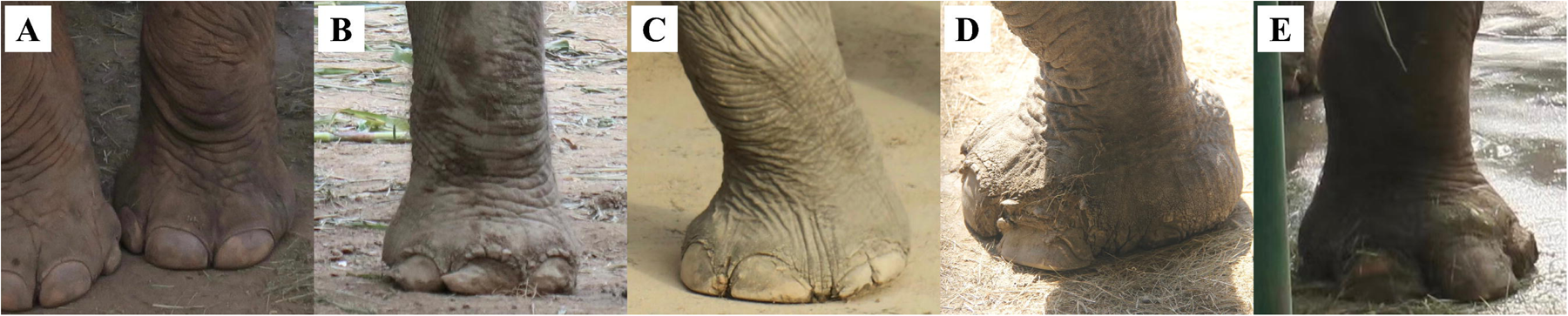
Samples of visible foot disorders of captive Asian elephants (Elephas maximus) recorded during on-site investigation. (A) feet in healthy appearance; (B) overgrown nails; (C) nail cracks; (D)overgrown cuticles; (E) joint deformation.

Here we hypothesized that both a larger outdoor enclosure area and a longer outdoor time could associate with lower BCS in an elephant while having foot disorders might associate with higher BCS.

### Statistical analysis

All statistical analyses were conducted using R version 4.2.2. In all calculations, sex was one of the categorical variables, with males and females encoded as 1 and 0 respectively. Individuals with unknown ages were excluded from the dataset.

To explore the relationship between BCS and its potential relative factors, we employed the package ‘prcomp’ to perform a principal component analysis (PCA) including seven independent variables (sex, age, daily feed supply, proportion of high-calorie feed, outdoor enclosure area, outdoor time, and foot disorder) and BCS. Calculated component scores and loadings were visualized using a biplot.

For further analysis, we employed the package ‘lmer’ to construct a stepwise multivariable linear mixed model (LMM) with the eight variables (sex, age, facility category, daily feed supply, proportion of high-calorie feed, outdoor enclosure area, outdoor time, and foot disorder) as fixed effects and different facilities as random effects. The optimal model selection was determined using the Akaike information (AIC) criterion. Next, we examined the interaction effects between variables in the optimal model and derived the optimal interaction model. Finally, in conjunction with significance tests, we assessed the significance of the relationships between BCS and its potential relative factors.

## Results

### Overview of captive Asian elephants’ body condition in China

We evaluated the BCS of 204 captive Asian elephants from 43 facilities in China (**Table 3**), and it suggested that 72.55% of elephants were in obesity to various degrees (BCS ≥ 7). Compared to elephants in the Wild Elephant Valley, elephants in zoos had significantly higher BCS (Independent-Samples T-Test, p < 0.001) on average.

**Table 3.** Overview of captive Asian elephants’ body condition score (BCS) in China.

| **Facility** | **Sex** | **BCS**<br>**Mean ± SE** | **BCS**<br>**Median** | **Proportion of**<br>**BCS ≥ 7** |
| --- | --- | --- | --- | --- |
| All | Both<br>(n=204) | 7.07±0.007 | 7 | 72.55% |
| | Male<br>(n=88) | 6.84±0.017 | 7 | 69.32% |
| | Female<br>(n=116) | 7.24±0.010 | 7 | 75.00% |
| Zoo | Both<br>(n=170) | 7.43±0.006 | 7 | 82.35% |
| | Male<br>(n=73) | 7.33±0.013 | 7 | 80.82% |
| | Female<br>(n=97) | 7.51±0.010 | 7 | 83.51% |
| Wild Elephant<br>Valley | Both<br>(n=34) | 5.26±0.044 | 5.5 | 23.53% |
| | Male<br>(n=15) | 4.47±0.100 | 4 | 13.33% |
| | Female<br>(n=19) | 5.89±0.061 | 6 | 31.58% |

### Relationship between BCS and its potential relative factors

The PCA analysis reduced the dataset from eight variables (seven independent variables and BCS) to two principal components. The first principal component (PC1) and the second principal component (PC2) accounted for 38.73% and 16.76% of the variance respectively. Therefore, the biplot of PC1 and PC2 represented more than half of the dataset’s variance (**Figure 3**). It suggested a positive correlation between BCS and the proportion of high-calorie feed. In contrast, there were strong negative correlations between the outdoor enclosure area and BCS, as well as between the outdoor time and BCS.

**Figure 3.**
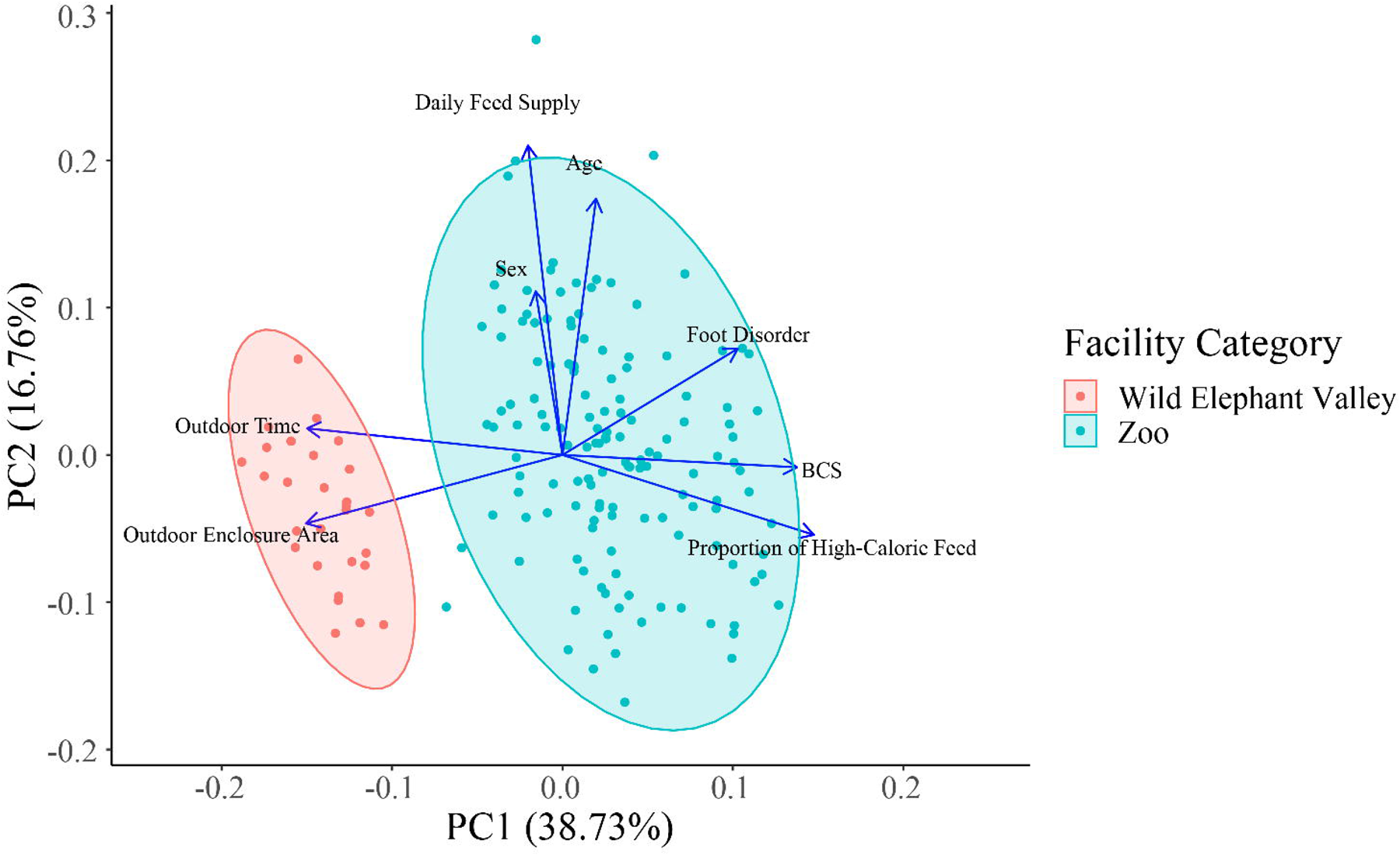
Biplot of principal component analysis (PCA) of BCS and its potential relative factors for all individuals of known age (n = 173). Points in the plot represented individual elephants, and eigenvectors show relations between the variables. Eigenvectors pointing in similar directions are positively correlated in the first two principal components, whereas eigenvectors pointing in opposite directions are negatively correlated. The length of each eigenvector represents its contribution to the corresponding variable.

For all individuals of known age (n = 173), the optimal linear model selected by stepwise regression included three BCS predictor variables: sex (p < 0.01), outdoor enclosure area (p < 0.001), and outdoor time (p < 0.001) (**Table S1**). ANOVA revealed that the BCS regression models with the interaction of sex and outdoor enclosure area, as well as outdoor time as independent variables, or the interaction of sex and outdoor time and outdoor enclosure area as independent variables, were statistically equivalent (**Table S2**). Considering that facilities generally do not keep any sex of elephants indoors during suitable weather conditions, we constructed a mixed regression model with BCS as the dependent variable, including four BCS predictor variables: (i) sex; (ii) outdoor enclosure area; (iii) outdoor time; and (iv) the interaction between sex and outdoor enclosure area (‘sex’×‘outdoor enclosure area’) as the fixed effects, whereas different facilities as the random effects. The results suggested that both outdoor enclosure area and outdoor time, along with the interaction between sex and outdoor enclosure area, were significantly negatively correlated with BCS (p < 0.05) (**Table 4**), indicating that both larger outdoor enclosure area and longer outdoor time are associated with lower BCS.

**Table 4.** Significance test of BCS predictor variables for all individuals of known age (n = 173)

| **Fixed Effects** | **Estimate** | **SE** | **df** | **t-value** | **p-value (>\|t\|)** |
| --- | --- | --- | --- | --- | --- |
| (Intercept) | 9.373e+00 | 3.697e-01 | 1.139e+02 | 25.350 | < 2e-16 *** |
| Sex | -1.034e-01 | 1.822e-01 | 4.780e+01 | -0.567 | 0.57311 |
| Outdoor Enclosure Area | -1.378e-04 | 4.868e-05 | 1.572e+02 | -2.831 | 0.00524 ** |
| Outdoor Time | -2.133e-01 | 4.296e-02 | 9.380e+01 | -4.964 | 3.08e-06 *** |
| ‘Sex’×‘Outdoor Enclosure Area’ | -2.540e-04 | 8.103e-05 | 4.838e+00 | -3.134 | 0.02702 * |

-Significance Code: ‘\*\*\*’ ~ p < 0.001; ‘\*\*’ ~ p < 0.01; ‘\*’ ~ p < 0.05

Taking into account the considerable disparity in outdoor enclosure areas between elephants in the Wild Elephant Valley and those of elephants in the zoo, we employed the same approach to select BCS predictors for individuals of known age in zoos (n = 144) (**Table S3**). The results indicated that only outdoor time exhibited a significant negative correlation with BCS (p < 0.01) (**Table 5**).

**Table 5.** Significance test of BCS predictor variables of BCS for individuals of known age in zoos (n = 144)

| **Fixed Effects** | **Estimate** | **SE** | **df** | **t-value** | **p-value (>\|t\|)** |
| --- | --- | --- | --- | --- | --- |
| (Intercept) | 9.01193 | 0.45191 | 24.58927 | 19.942 | < 2e-16 *** |
| Outdoor Time | -0.18904 | 0.05397 | 22.61886 | -3.503 | 0.00195 ** |

-Significance Code: ‘\*\*\*’ ~ p < 0.001; ‘\*\*’ ~ p < 0.01

## Discussion

In this study, we examined the relationship between potential related factors and the obesity status of captive Asian elephants in China by body condition assessment and linear regression. Study results indicated that longer outdoor time was related to lower BCS of an elephant. Here we conducted the following discussion for various factors associated with diet and exercise, as well as situations of elephants in different facilities.

### Enclosures, exhibition, and obesity of captive elephants in zoos

Linear regression showed that for all individuals of known age, both outdoor enclosure area and outdoor time were significantly negatively correlated with BCS. However, for elephants in zoos, no significant correlation between outdoor enclosure area and BCS was observed. One of the possible reasons for this is that outdoor enclosures provided by zoos for elephants are generally too small in China.

For welfare considerations, facilities should provide outdoor enclosures as large as possible and enrichment for elephants, which may encourage elephants to display natural behaviors and increase their amount of exercise to alleviate or avoid obesity (Hacker *et al* 2018). A study on captive Asian elephants in European zoos showed that females in smaller enclosures are more prone to obesity (Schiffmann *et al* 2018). Although CAZG has not imposed strict requirements on outdoor enclosure areas for captive elephants, it has recommended that the outdoor enclosure area should not be less than 170 m^2^ for each elephant on average (Zhang *et al* 2018). Despite these guidelines, our investigation revealed that nearly half (48.82%) of elephants in zoos still did not reach CAZG’s recommended standard. Since it is difficult for most zoos to expand outdoor enclosures in the short term, we suggest that it is crucial and practical for Chinese zoos to improve the utilization rate of limited outdoor enclosures for elephants through diverse enrichment, for instance, pools, sand pits, wallows, stumps for tickling, shade canopies, and foraging devices, etc. Moreover, it is also notable to maintain a suitable population size of elephants for a zoo. Since elephants have intricate social structures and emotions, housing only one elephant in captivity could have detrimental effects on its mental health over time. Conversely, an overcrowded population (e.g., housing over ten elephants in the same enclosure) could lead to cramped living space for each elephant.

Multiple studies indicated that inactivity and long-term indoor living significantly increase the risk of degenerative lesions in the musculoskeletal system of captive elephants, aggravating obesity and even leading to disability or euthanasia (Greco *et al* 2016; Miller *et al* 2016; Bansiddhi *et al* 2020). For the relationship between outdoor time and the obesity status of captive elephants, a study on captive Asian elephants in North American zoos has shown that staff-directed walking exercise of 14 hours or more per week can effectively reduce the risk of obesity (Morfeld *et al* 2016). However, another study based on a 10-month observation of captive Asian elephants in a British zoo revealed that among the 6.5-hour outdoor exhibition every day, adult elephants spent most of their outdoor time feeding, standing still, or resting, and spent only 6.1-19.2% of outdoor time walking (Rees 2009). Here we suggest that staff members could take measures to extend the outdoor time of captive elephants, for instance, providing elephants with options to freely enter and exit indoors during closing hours of zoos. Additionally, facilities may install outdoor heat sources to extend the outdoor time of elephants in lower environmental temperatures.

### Diet and obesity of captive elephants

Although statistical analysis showed that both daily feed supply and the proportion of high-calorie feed were insignificantly related to BCS, since we could not measure the exact feed intakes of elephants in this study, further research is required to determine the effects of diet on obesity of captive elephants in China. It is recommended that the cellulose content in captive elephant feed should be no less than 30%, and herb forage should be no less than 70% of the total mass of daily feed to avoid indigestion (Ullrey *et al* 1997; Romain *et al* 2014). The supply of high-calorie feed (fruits, vegetables, pellets, etc.) should be controlled with proper supplementation of required trace elements based on specific situations to avoid obesity caused by overeating and intestinal gas caused by excessive protein intake (Hatt & Clauss 2006). However, our investigation revealed that the proportion of high-calorie feed was over 30% in the diet of over a quarter (27.45%) of elephants. Furthermore, some zoos used to add starchy foods such as pumpkins and sweet potatoes to their elephant feed in winter. Considering that elephants stay indoors all day during this period, it might be related to the higher BCS because of reduced exercise and increased nutrient intake. Here we suggest that staff members could control the supply of high-calorie feed, use feeding devices as enrichment, and try temporally unpredictable feeding schedules, which might contribute to the body weight management of captive elephants (Holdgate *et al* 2016; Scott & LaDue 2019).

### Foot health of captive Asian elephants in China

Although statistical analysis showed that foot disorder was insignificantly related to BCS, our investigation indicated that 58.24% of zoo elephants had foot disorders of varying degrees. A study in European zoos revealed that time spent indoors and on hard surfaces was positively correlated with the risk of foot disorders, while natural, soft ground (e.g., sand flooring) could alleviate this situation (Wendler *et al* 2019). However, all 42 investigated zoos used hard cement floors in their indoor spaces. To guarantee the health of feet, European and North American zoos provide their elephants with regular foot health checks based on protected contact and positive reinforcement training (Laule & Whittaker 2009). Unfortunately, less than 25% of the 42 zoos have corresponding devices (such as erected training walls) and relevant technical personnel for regular foot health checks. Moreover, many hidden foot lesions need to be diagnosed by CT scanning or the elephant’s posture during locomotion instead of simply observing the appearance at the initial stage (Panagiotopoulou *et al* 2016; Regnault *et al* 2017). It implied that the foot health problems of captive elephants in China might be much more serious than observed. To improve this situation, we suggest facilities enhance staff members’ awareness of the importance of foot health for captive elephants. Moreover, it is also necessary to promote positive reinforcement training in captive elephants for regular foot health checks.

### Elephants in the Wild Elephant Valley

According to our investigation and statistical analysis, there were significantly lower BCS on average and no visible foot disorders for elephants in the Wild Elephant Valley, which might be attributed to local environmental and husbandry conditions similar to wild Asian elephant habitats. Nevertheless, we suggested that long-term engagement in performance might harm the mental health of elephants, leading to stereotypical behavior. Furthermore, since captive elephants in the Wild Elephant Valley were not completely isolated from local wild elephant populations, some pathogens (e.g., EEHVs) carried by captive elephants might spread during their roaming in natural forests and cause the infection of wild individuals (Yang *et al* 2022).

### Animal welfare implications

This study was the first relevant research on the obesity status and foot health of captive Asian Elephants in China. Honestly, there were still a series of limitations in our study. For instance, our investigation did not include all captive Asian elephants in China. Restricted by the veterinary conditions of elephant-raising facilities, we could not obtain tissue samples for further biochemical metabolic examination of obesity. Although body condition assessment may not be able to examine the obesity status of elephants more accurately than direct weighing and measuring serum leptin (Chusyd *et al* 2021), we hoped that our study could arouse the attention of relevant facilities on the welfare problems, including obesity, of captive Asian elephants in China. We also hoped that our work could inspire the future husbandry management of captive elephants in China and provide a reference for follow-up studies on welfare issues in captive megaherbivores.

## Conclusion

Our study indicated that insufficient outdoor time might be the primary potential cause of the prevalent high BCS of captive Asian elephants in China. Here we suggested elephant-raising facilities to take measures to extend the outdoor time of elephants. Moreover, a suitable diet with feeding enrichment and regular foot health checks might also contribute to the body weight management of captive elephants.

## Supporting information

dataset for statistical analysis

operation process of R

## Acknowledgements

Thanks for the funding support and permission to work with each elephant-raising facility from CAZG. We were also very grateful for the cooperation of all 43 elephant-raising facilities. As the first nationwide welfare-related study of captive elephants in China, we sincerely hoped to keep this cooperative relationship and work together to improve the welfare of captive elephants in the future.

