## Supplementary material for "Obesity status and its relative factors of captive Asian elephants (*Elephas maximus*) in China based on body condition assessment": Table S1 to S3

**Table S1 Selection of BCS predictor variables by** **stepwise regression for** **all individuals of known age (n = 173)**

|  | **Variable** | **Eliminated** | **Number of Parameters in Reduced Models**  **(npar)** | **Log-likelihood of the Given Model**  **(logLik)** | **AIC** | **LRT** | **df** | **Pr (>χ^2^)** |
| --- | --- | --- | --- | --- | --- | --- | --- | --- |
| **Random Effect** | <none> |  | 11 | -268.24 | 558.48 |  |  |  |
|  | Facility Number | 1 | 10 | -268.98 | 577.96 | 1.4752 | 1 | 0.2245 |
|  | **Variable** | **Eliminated** | **df** | **Sum of Squares** | **RSS** | **AIC** | **F-value** | **Pr (>F)** |
| **Fixed Effect** | Facility Category | 1 | 1 | 0.009 | 165.96 | 8.814 | 0.0084 | 0.927001 |
|  | Age | 2 | 1 | 0.203 | 166.16 | 7.025 | 0.2017 | 0.653922 |
|  | Foot Disorder | 3 | 1 | 1.167 | 167.33 | 6.236 | 1.1658 | 0.281831 |
|  | Daily Feed Supply | 4 | 1 | 2.168 | 169.50 | 6.464 | 2.1641 | 0.143150 |
|  | Proportion of High-Calorie Feed | 5 | 1 | 1.211 | 170.71 | 5.696 | 1.2007 | 0.274743 |
|  | Sex | 0 | 1 | 8.976 | 179.69 | 12.561 | 8.8862 | 0.003298 ** |
|  | Outdoor Enclosure Area | 0 | 1 | 32.759 | 203.47 | 34.065 | 32.4309 | 5.362e-08 *** |
|  | Outdoor Time | 0 | 1 | 22.708 | 193.42 | 25.301 | 22.4808 | 4.487e-06 *** |

-Significance Code: ‘***’ ~ p < 0.001; ‘**’ ~ p < 0.01

**Table S2 Selection of optimal interaction model of BCS predictor variables for all individuals of known age (n = 173)**

|  | **Number of Parameters in Reduced Models (npar)** | **AIC** | **BIC** | **Deviance of Log-likelihood of the Given Model**  **(logLik Deviance)** | **χ^2^** | **df** | **Pr (>χ^2^)** |
| --- | --- | --- | --- | --- | --- | --- | --- |
| lmer.fit2 | 7 | 487.47 | 500.54 | -236.73 | 473.47 |  |  |
| lmer.fit3 | 7 | 499.66 | 521.73 | -242.83 | 485.66 | 0 | 0 |

\# lmer.fit2 ← lmer (BCS ~ ‘Sex’×‘Outdoor Enclosure Area’ + ‘Outdoor Time’ + (1 |‘Facility Number’),

\# lmer.fit3 ← lmer (BCS ~ ‘Sex’×‘Outdoor Time’ + ‘Outdoor Enclosure Area’ + (1 |‘Facility Number’),

**Table S3 Selection of BCS predictor variables by stepwise regression for individuals of known age in zoos (n = 144)**

|  | **Variable** | **Eliminated** | **Number of Parameters in Reduced Models**  **(npar)** | **Log-likelihood of the Given Model**  **(logLik)** | **AIC** | **LRT** | **df** | **Pr (>χ^2^)** |
| --- | --- | --- | --- | --- | --- | --- | --- | --- |
| **Random Effect** | <none> |  | 10 | -210.54 | 441.08 |  |  |  |
|  | Facility Number | 0 | 9 | -212.80 | 443.60 | 4.5203 | 1 | 0.0335 * |
|  | **Variable** | **Eliminated** | **Sum of Squares** | **Mean Square** | **Numerator Degrees of Freedom (NumDF)** | **Denominator Degrees of Freedom**  **(DenDF)** | **F-value** | **Pr (>F)** |
| **Fixed Effect** | Age | 1 | 0.1648 | 0.1648 | 1 | 135.935 | 0.2406 | 0.62457 |
|  | Foot Disorder | 2 | 0.1678 | 0.1678 | 1 | 70.361 | 0.2475 | 0.62038 |
|  | Outdoor Enclosure Area | 3 | 0.8358 | 0.8358 | 1 | 61.793 | 1.2391 | 0.26996 |
|  | Daily Feed Supply | 4 | 1.0280 | 1.0280 | 1 | 54.429 | 1.5345 | 0.22075 |
|  | Proportion of High-Calorie Feed | 5 | 0.7006 | 0.7006 | 1 | 35.520 | 1.0421 | 0.31424 |
|  | Sex | 6 | 1.8350 | 1.8350 | 1 | 121.531 | 2.7363 | 0.10067 |
|  | Outdoor Time | 0 | 8.3285 | 8.3285 | 1 | 22.619 | 12.2692 | 0.00195 ** |

-Significance Code: ‘**’ ~ p < 0.01; ‘*’ ~ p < 0.05
