## Supplementary material for "Obesity status and its relative factors of captive Asian elephants (*Elephas maximus*) in China based on body condition assessment": operation process of R

### Analysis outputs of R

#### Resubmission of Manuscript Z2647

##### PCA

```r
dat <- read.csv('~/Downloads/data_AW_submission.csv', check.names = F) %>% drop_na()
pca.fit <- prcomp(dat[,3:10], scale. = T)
fviz_eig(pca.fit, addlabels = T)
```

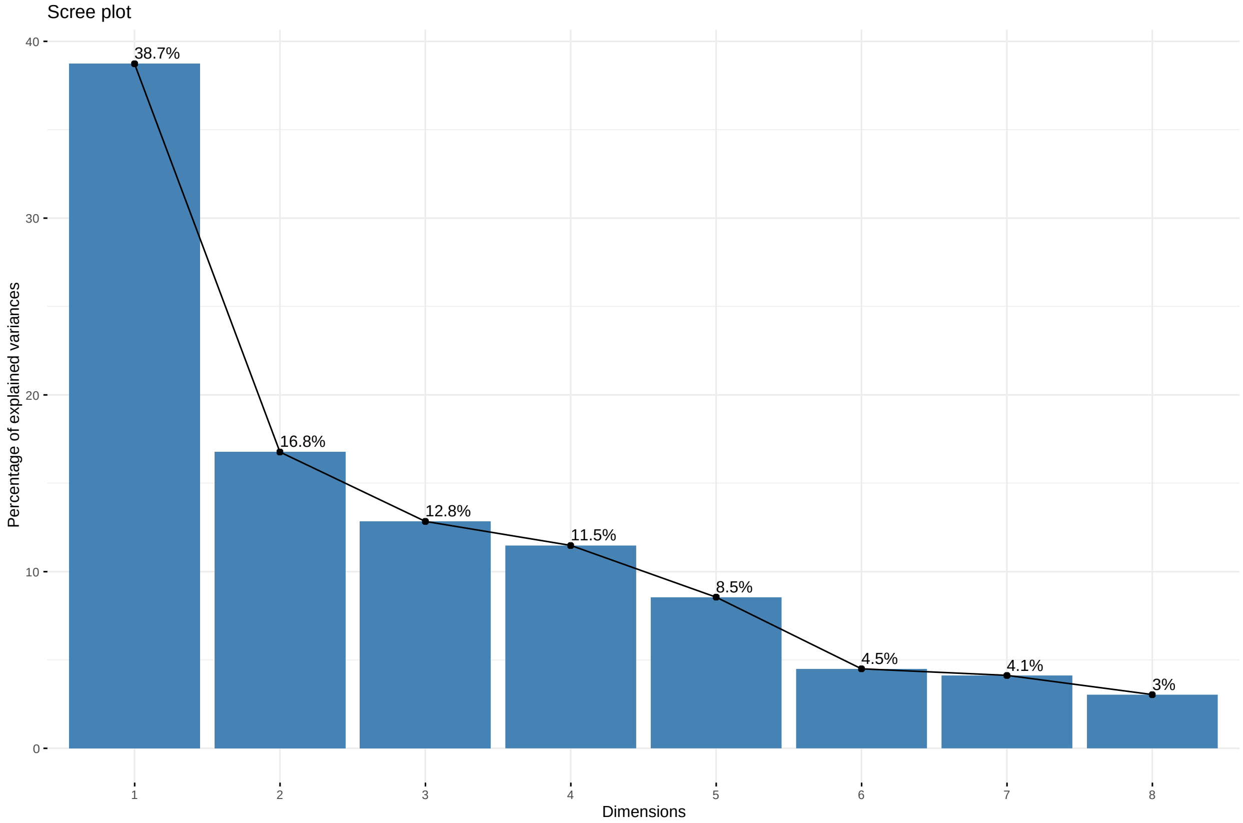


```r
autoplot(pca.fit, data = dat, colour = 'Facility_Category', loadings = T,
              loadings.colour = 'blue', loadings.size = 1,
  frame = TRUE, frame.type = 'norm') +
  theme_classic() +
  labs(fill = 'Facility Category', color = 'Facility Category') +
  xlim(c(-0.24, 0.25)) +
  annotate("text", x = (pca.fit$rotation[,1]/2.7-0.01), y = (pca.fit$rotation[,2]/2.8),
      label = rownames(pca.fit$rotation)) +
  theme(text = element_text(size = 20))
```

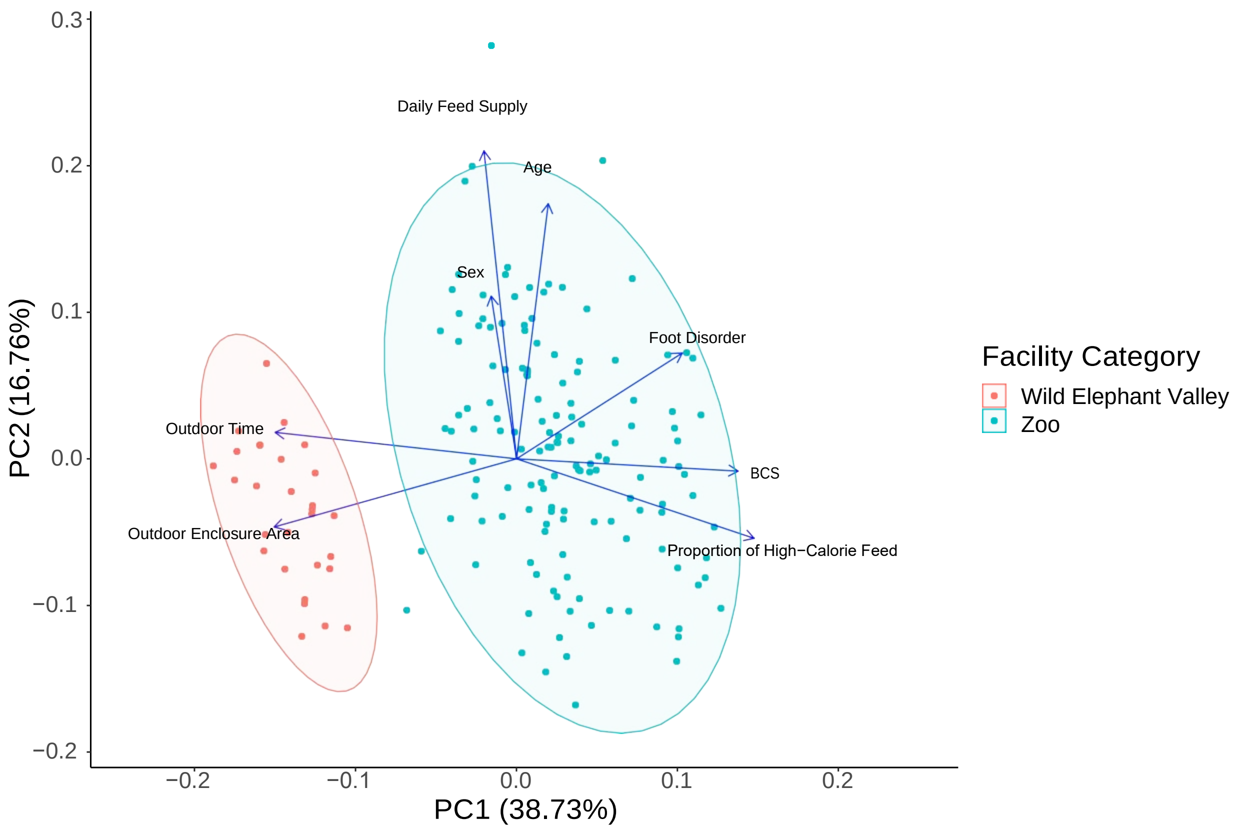


#### LMM

##### Variable selection

```r
dat$Sex <- as.factor(dat$Sex)
dat$`Foot Disorder` <- as.factor(dat$`Foot Disorder`)
lmer.fit <- lmer(BCS ~ Sex + Facility_Category + Age + `Foot Disorder` +
                   `Daily Feed Supply` + `Proportion of High-Calorie Feed` +
                   `Outdoor Enclosure Area` + `Outdoor Time` +
                   (1|`Facility Number`),
                 data = dat)
step(lmer.fit)
```

```
## Backward reduced random-effect table: ##

##                          Eliminated npar logLik   AIC    LRT Df Pr(>Chisq)
## <none>                              11 -268.24 558.48
## (1 | 'Facility Number')           1 10 -268.98 557.96 1.4752 1     0.2245
##
## Backward reduced fixed-effect table: ##
## Facility_Category        Eliminated Df Sum of Sq    RSS   AIC F value
## Age                               1  1     0.009 165.96 8.814  0.0084
## 'Foot Disorder'                   2  1     0.203 166.16 7.025  0.2017
                                     3  1     1.167 167.33 6.236  1.1658
```

```
## ‘Daily Feed Supply‘                    4  1    2.168 169.50   6.464   2.1641
## ‘Proportion of High-Calorie Feed‘      5  1    1.211 170.71   5.696   1.2007
## Sex                                    0  1    8.976 179.69  12.561   8.8862
## ‘Outdoor Enclosure Area‘               0  1   32.759 203.47  34.065  32.4309
## ‘Outdoor Time‘                         0  1   22.708 193.42  25.301  22.4808
##                                  Pr(>F)
## Facility_Category               0.927001
## Age                             0.653922
## ‘Foot Disorder‘                 0.281831
## ‘Daily Feed Supply‘             0.143150
## ‘Proportion of High-Calorie Feed‘ 0.274743
## Sex                             0.003298 **
## ‘Outdoor Enclosure              5.362e-08 ***
Area‘ ## ‘Outdoor Time‘         4.487e-06 ***
## ---
## Signif. codes: 0 ’***’ 0.001 ’**’ 0.01 ’*’ 0.05 ’.’ 0.1 ’ ’ 1 ##
## Model found:
## BCS ~ Sex + ‘Outdoor Enclosure Area‘ + ‘Outdoor Time‘
```

**Interaction model**

```
lmer.fit2 <- lmer(BCS ~ Sex * `Outdoor Enclosure Area` + `Outdoor Time` +
                    (1 |`Facility Number`),
                  data = dat)
lmer.fit3 <- lmer(BCS ~ Sex * `Outdoor Time` + `Outdoor Enclosure Area` +
                    (1 |`Facility Number`),
                  data = dat)
anova(lmer.fit2, lmer.fit3)
```

```
## refitting model(s) with ML (instead of REML)

## Data: dat
## Models:
## lmer.fit2: BCS ~ Sex * ‘Outdoor Enclosure Area‘ + ‘Outdoor Time‘ + (1 | ‘Facility Number‘) ## lmer.fit3:
BCS ~ Sex * ‘Outdoor Time‘ + ‘Outdoor Enclosure Area‘ + (1 | ‘Facility Number‘)
##           npar    AIC    BIC  logLik deviance Chisq Df Pr(>Chisq)
## lmer.fit2    7 487.47 509.54 -236.73   473.47
## lmer.fit3    7 499.66 521.73 -242.83   485.66     0  0
```

```
lmer.fit <- lmer(BCS ~ Sex * `Outdoor Enclosure Area` + `Outdoor Time` +
                   (1 |`Facility Number`),
                 data = dat)
```

```
summary(lmer.fit)
```

```
## Linear mixed model fit by REML. t-tests use Satterthwaite's method [ ##
lmerModLmerTest]
## Formula: BCS ~ Sex * ‘Outdoor Enclosure Area‘ + ‘Outdoor Time‘ + (1 |
##         ‘Facility Number‘)
##       Data: dat
##
## REML criterion at convergence: 518.2
##
## Scaled residuals:
##          Min            1Q      Median            3Q          Max
##  -2.45260  -0.75607    0.05398    0.53576    3.02309
##
## Random effects:
##   Groups                 Name            Variance  Std.Dev.
##    Facility  Number (Intercept)  0.09501     0.3082
##   Residual                               0.86089    0.9278
##  Number  of  obs: 173,  groups:    Facility Number, 42 ##
## Fixed effects:

##                                              Estimate  Std.  Error                  df t value Pr(>|t|)
## (Intercept)                                9.220e+00    4.242e-01    2.597e+01    21.734    <  2e-16
## Sex1                                      −1.211e-01    1.710e-01    1.610e+02    -0.708  0.479960
## ‘Outdoor Enclosure                        −1.460e-04    7.194e-05    6.634e+00    -2.029  0.084209
Area‘ ## ‘Outdoor Time‘                   −1.943e-01    5.024e-02    2.150e+01    -3.867  0.000862
##  Sex1:‘Outdoor  Enclosure  Area‘  -2.550e-04       6.354e-05    1.329e+02    -4.014  9.94e-05
##
## (Intercept)                              ***
## Sex1
## ‘Outdoor Enclosure                       .
Area‘ ##  ‘Outdoor  Time‘                  ***
## Sex1:‘Outdoor Enclosure Area‘ ***
## ---
##  Signif.  codes:    0 ’***’ 0.001 ’**’ 0.01 ’*’ 0.05 ’.’ 0.1 ’ ’ 1 ##
## Correlation of Fixed Effects:

##                      (Intr)  Sex1      ‘OEAr‘  ‘OtTm‘
##  Sex1                 -0.212
##  ‘OtdrExcAr‘    0.296    0.146
##  ‘OutdoorTm‘  -0.954    0.042  -0.433
##  Sx1:‘OtEAr‘    0.101 -0.521 -0.354 -0.017
```

```
shapiro.test(resid(lmer.fit))
```

```
##
##    Shapiro-Wilk normality test ##
##  data:    resid(lmer.fit)
## W = 0.99187, p-value = 0.4394
```

```
qqnorm(resid(lmer.fit))
qqline(resid(lmer.fit))
```

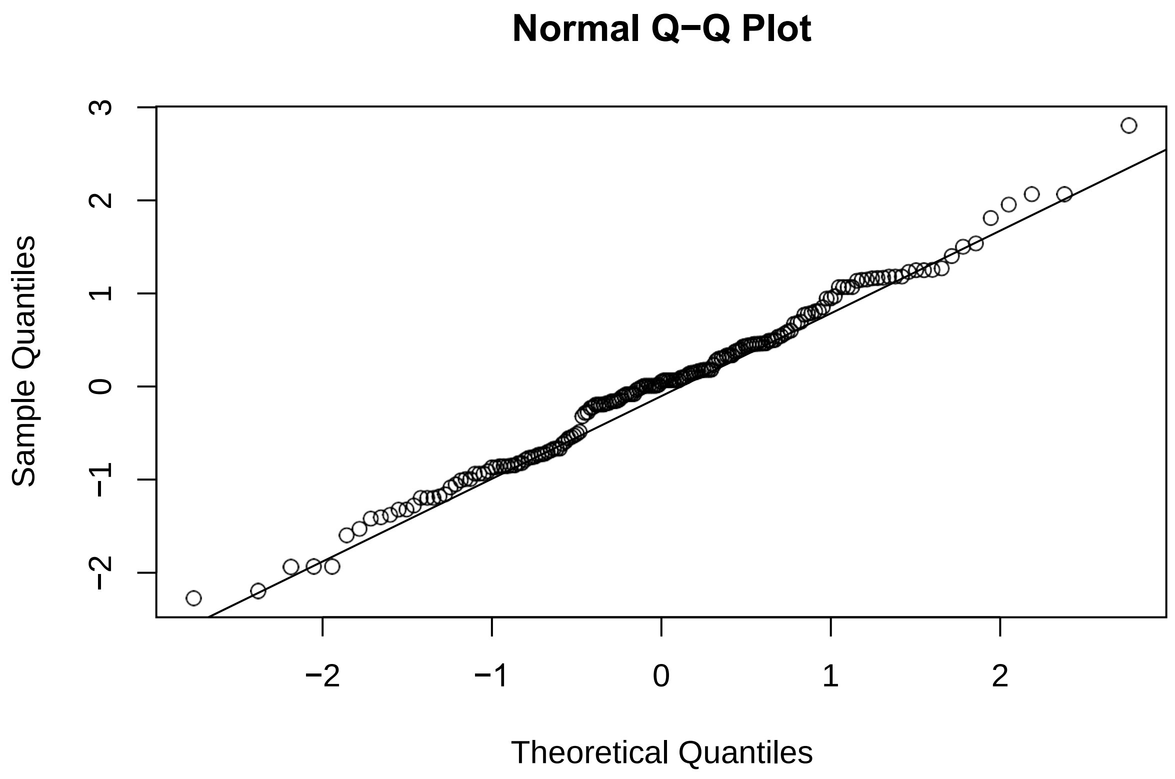


```
ggpredict(lmer.fit, terms = c('Outdoor Time')) %>%
  plot(add.data =T, ci =F) +
  theme_classic()
```

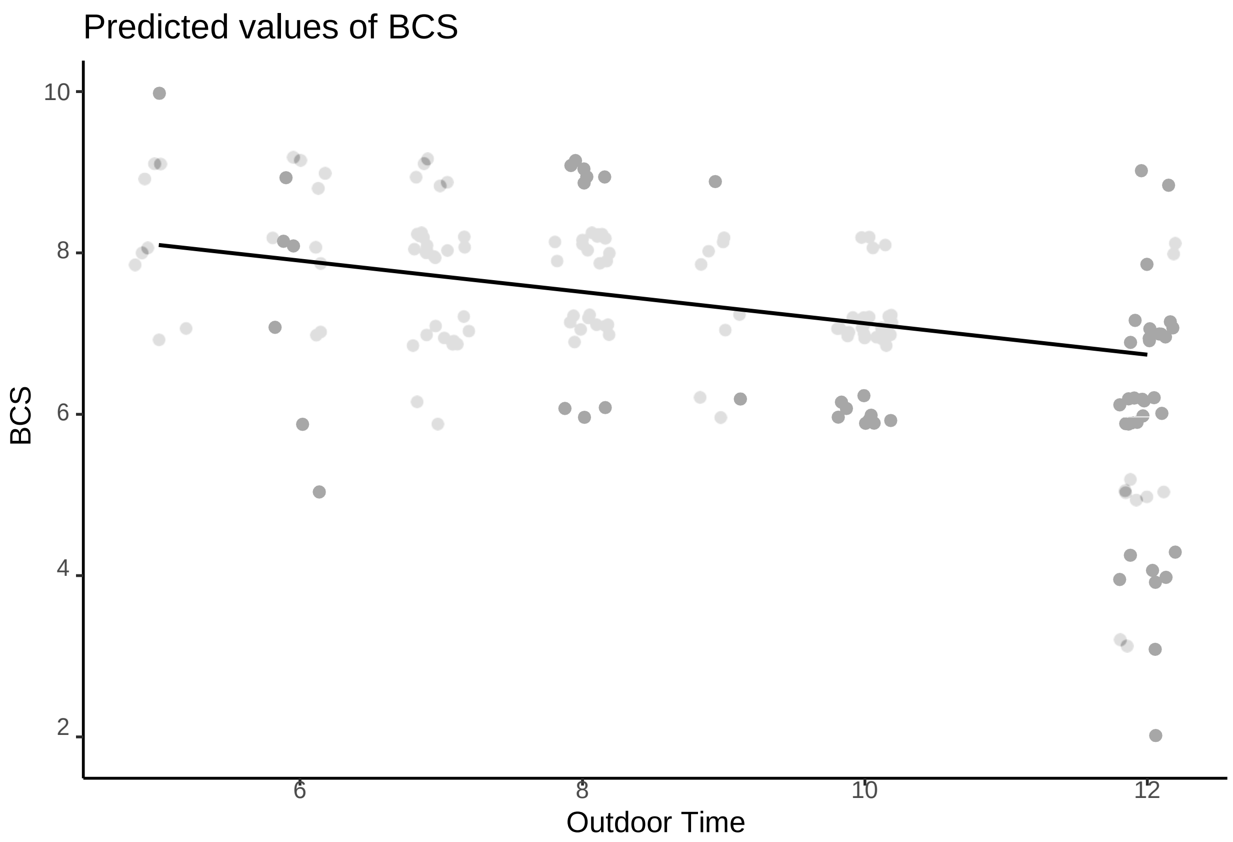


```
ggpredict(lmer.fit, terms = c('Outdoor Enclosure Area', 'Sex')) %>%
  plot(add.data =T, ci =F) +
  theme_classic()
```

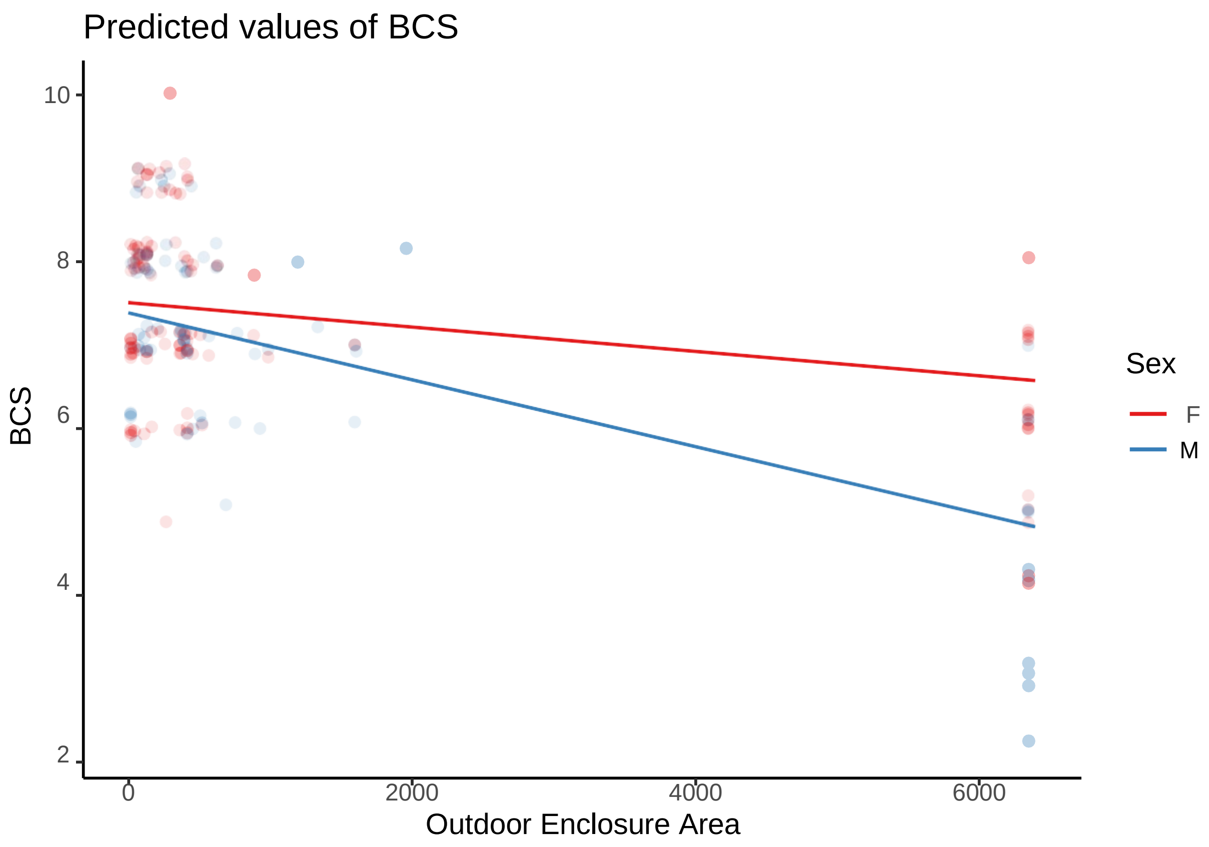

Predicted values of BCS
10
8
6
4
2
BCS
0
2000
4000
6000
Outdoor Enclosure Area
Sex
F
M

**LMM for zoo data**

**Variable selection**

```
dat$Sex <- as.factor(dat$Sex)
dat$`Foot Disorder` <- as.factor(dat$`Foot Disorder`) lmer.fit
<- lmer(BCS ~ Sex + Age + `Foot Disorder` +
                    `Daily Feed Supply` + `Proportion of High-Calorie Feed` +
                    `Outdoor Enclosure Area` + `Outdoor Time` +
                    (1|`Facility Number`),
                    data = dat %>% filter(Facility_Category = = 'Zoo'))
step(lmer.fit)
```

```
## Backward reduced random-effect table: ##

##                                Eliminated  npar   logLik      AIC     LRT Df Pr(>Chisq)
## <none>                                       10 -210.54  441.08
##  (1  | 'Facility  Number')              0       9  -212.80  443.60  4.5203    1       0.0335  *
## ---
##  Signif.  codes:   0  '***'  0.001  '**'  0.01  '*'  0.05  '.'  0.1  ' '  1 ##
## Backward reduced fixed-effect table:
## Degrees of freedom method: Satterthwaite
##

##                                        Eliminated Sum Sq Mean Sq NumDF DenDF
##  Age                                            1 0.1648    0.1648       1  135.935
##  'Foot Disorder'                                2 0.1678    0.1678       1   70.361
##  'Outdoor Enclosure Area'                       3 0.8358    0.8358       1   61.793
##  'Daily Feed Supply'                            4 1.0280    1.0280       1   54.429
##  'Proportion of High-Calorie    Feed' ##        5 0.7006    0.7006       1   35.520
Sex                                             6 1.8350    1.8350       1  121.531
##  'Outdoor Time'                                 0 8.3285    8.3285       1   22.619
##                                        F  value    Pr(>F)
##  Age                                     0.2406 0.62457
##  'Foot Disorder'                         0.2475 0.62038
##  'Outdoor Enclosure Area'                1.2391 0.26996
##  'Daily Feed                             1.5345 0.22075
##  Supply' 'Proportion of High-  Feed'   1.0421  0.31424
     Calorie
##  Sex                                     2.7363 0.10067
##   Outdoor Time'                         12.2692 0.00195 **
##  ---
##  Signif.  codes:   0  '***'  0.001  '**'  0.01  '*'  0.05  '.'  0.1  ' '  1 ##
## Model found:
## BCS ~ 'Outdoor Time' + (1 | 'Facility Number')
```

```
lmer.fit <- lmer(BCS ~ `Outdoor Time` +
                   (1 |`Facility Number`),
                 data = dat %>% filter(Facility_Category = = 'Zoo'))
summary(lmer.fit)
```

```
## Linear mixed model fit by REML. t-tests use Satterthwaite's method [ ##
lmerModLmerTest]
## Formula: BCS ~ ‘Outdoor Time‘ + (1 | ‘Facility Number‘)
##       Data: dat %>% filter(Facility_Category == "Zoo")
##
## REML criterion at convergence: 383.6
##
## Scaled residuals:
##        Min          1Q    Median          3Q         Max
## -2.4362 -0.8197    0.1232    0.4298    2.5720
##
## Random effects:
##   Groups                  Name            Variance  Std.Dev.
##    Facility  Number (Intercept)  0.1960      0.4427
##   Residual                                 0.6788      0.8239
##  Number  of  obs: 144,  groups:    Facility Number, 41 ##
## Fixed effects:

##                          Estimate  Std.  Error               df t value  Pr(>|t|)
##  (Intercept)            9.01193       0.45191  24.58927    19.942    < 2e-16  ***
## ‘Outdoor Time‘ -0.18904           0.05397  22.61886    -3.503    0.00195  **
## ---
##  Signif.  codes:    0  ’***’  0.001  ’**’  0.01  ’*’  0.05  ’.’  0.1  ’  ’  1 ##
## Correlation of Fixed Effects:
##                     (Intr)
## ‘OutdoorTm‘ -0.973
```

```r
shapiro.test(resid(lmer.fit))
```

```
##
##    Shapiro-Wilk normality test ##
##  data:    resid(lmer.fit)
## W = 0.97888, p-value = 0.02531
```

```r
qqnorm(resid(lmer.fit))
qqline(resid(lmer.fit))
```

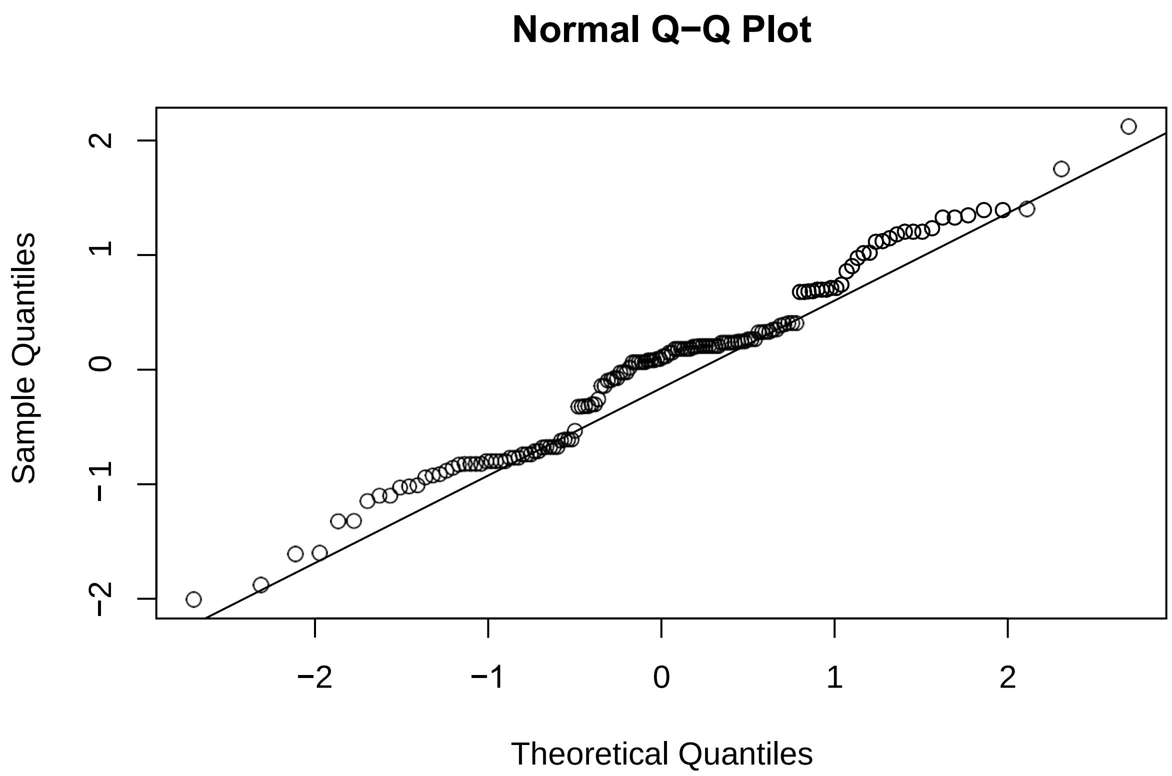


```r
ggpredict(lmer.fit, terms = c('Outdoor Time')) %>%
  plot(add.data =T, ci =F) +
  theme_classic()
```

Predicted values of BCS

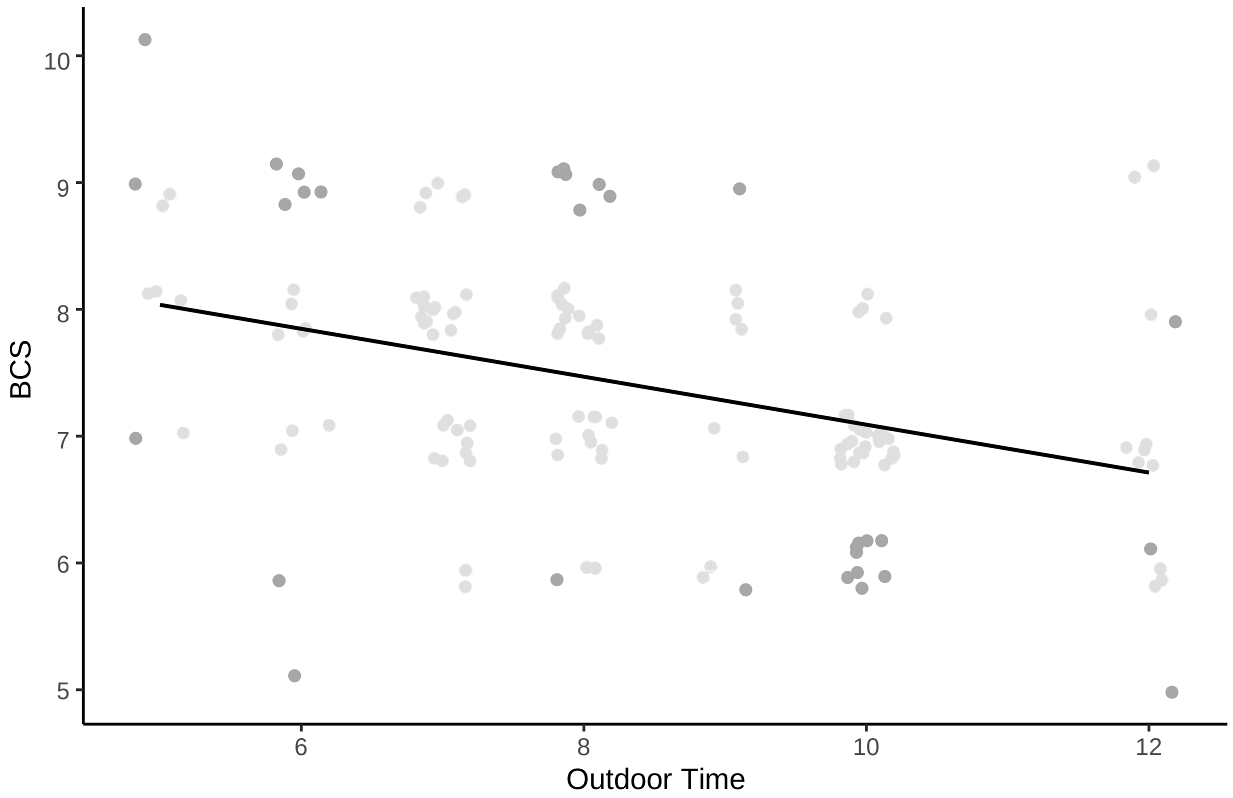
